# Shaping the Future: the importance of land use when projecting the future distribution of Endangered Amphibians and Reptiles

**DOI:** 10.1101/2024.07.25.605069

**Authors:** Nina Luisa Santostasi, Chiara Serafini, Luigi Maiorano

## Abstract

Obtaining reliable predictions of future species distributions under global change scenarios is fundamental for identifying priority conservation sites. This requires incorporating the main drivers of distribution shifts into the modeling process. Climate change and land use change are the two primary drivers of species distributions, yet they have been considered differently in studies exploring future scenarios. Most studies focus solely on bioclimatic variables, given that climate is a major determinant of species distributions on a large spatial scale.

In our study, we explored the impact of including dynamic land use variables in species distribution models, examining predicted range loss and spatial mismatch of suitable areas compared to models using only bioclimatic variables. Our findings indicate that including land cover can significantly alter model outputs. While incorporating land use and land cover variables did not enhance the predictive power for most species, it substantially affected the predicted future suitable areas for 60% of the species (mean change = 69%, range: 4% to 167%). This trend was consistent across all spatial resolutions.

The two modeling approaches also differed in the spatial location of suitable areas. The level of disagreement varied across species but was generally high for both current and future scenarios, increasing with coarser resolutions. Our results underscore the significant implications of excluding land use change variables and highlight the necessity of considering these factors on a taxon-specific basis.

## INTRODUCTION

In a context of global changes, species are often responding by shifting their distribution (Pecl et al. 2017). Obtaining reliable predictions of future species distribution under global change scenarios is therefore fundamental for the identification of priority sites for conservation (e.g., climatic refugia; Cavalcante et al., 2024; Serafini et al. submitted). Correlative species distribution models (SDMs) allow to predict environmental-driven range shifts by estimating a link between presence data and environmental variables and then forecast the species niche to future values of such variables (Elith et al., 2010).

Climate change and land use change are possibly the two major drivers of species distributions (Mantyka-pringle et al., 2012), but they have been considered differently in studies exploring future scenarios (Martin et al., 2013). The majority of these rely only on climate variables, because climate is considered the main determinant of species distributions at a large spatial scale (e.g., when considering regional to continental to global scales with spatial resolution > 10 km^2^; Thuiller et al., 2004; Bellard et al., 2012). Although modelling species distribution at large scales is appropriate to investigate biogeographical biodiversity patterns, finer scales (with resolution ≤ 10 km^2^) might be more appropriate for local conservation purposes. However, at local to regional scales the addition of land cover variables to pure bioclimatic models significantly improved the explanatory and predictive power of bioclimatic models for anurans, turtles and birds (Luoto et al., 2007, Tingley and Herman 2009). Most of these studies use static land cover variables because fine-scale dynamic land use future projections have become available only recently (e.g., Chen and Liu 2022). Therefore, the effect of considering them in fine scale SDMs and on their future projections implications remain largely unexplored.

To our knowledge only one study (Martin et al., 2013) examined the effect of using land use change scenarios along with climate change scenarios on future species distribution projections obtained by SDMs, compared with the classical approach (i.e., climate only models) and at different resolutions (50, 10 and 5 km^2^). This study concluded that even at the finest resolution, the inclusion of dynamic land use variables in species distribution did not significantly affect future projections suggesting that this was due to the poor thematic resolution of land use scenarios which included only 5 categories (built-up area, arable land, permanent crop, forest, grassland and others) that poorly represented habitat suitability for the species. Here, we further explore this issue taking advantage of the recently available land use change scenarios with improved thematic resolution (n = 9 thematic categories) and high spatial resolution (300 m^2^). We explored the impact of including dynamic land use variables in SDMs looking at predicted range loss and spatial mismatch of suitable areas when compared to bioclimatic only SDMs.

We focused our analyses on the Corsica-Sardinia microplate in the North-Western Mediterranean, including the two main islands (Corsica and Sardinia) plus their archipelagos (Fig: 1). Given their long history of isolation from the mainland (29 million years) and from each other (9 million years; Alvarez et al., 1972), the two islands are hotspots of endemism (Médail and Quèzel 1999). This insular system represents the ideal setting to develop fine-scale SDMs because the long-term geographic isolation and the presence of endemic species and/or populations allows to calibrate the models avoiding artificial niche truncation (Machado-Stredel et al., 2021). We focused endemic (present only in one or both islands) and sub-endemic (being also present in few other localities) herpetofauna (i.e., amphibians and reptiles) because these species are particularly sensitive to both climate and land use alterations. We developed climate only SDMs and climate + land use SDMs considering three different spatial resolutions, 10 km^2^, 5 km^2^, and 1 km^2^, and keeping the study area unchanged. We focused on the following questions: 1) does the importance of land use variables in SDMs change with spatial resolution? 2) does the predictive power change when comparing bioclimatic only SDMs with SDMs including also land cover variables? 3) do future predictions change when including land cover data?

**Fig. 1.**
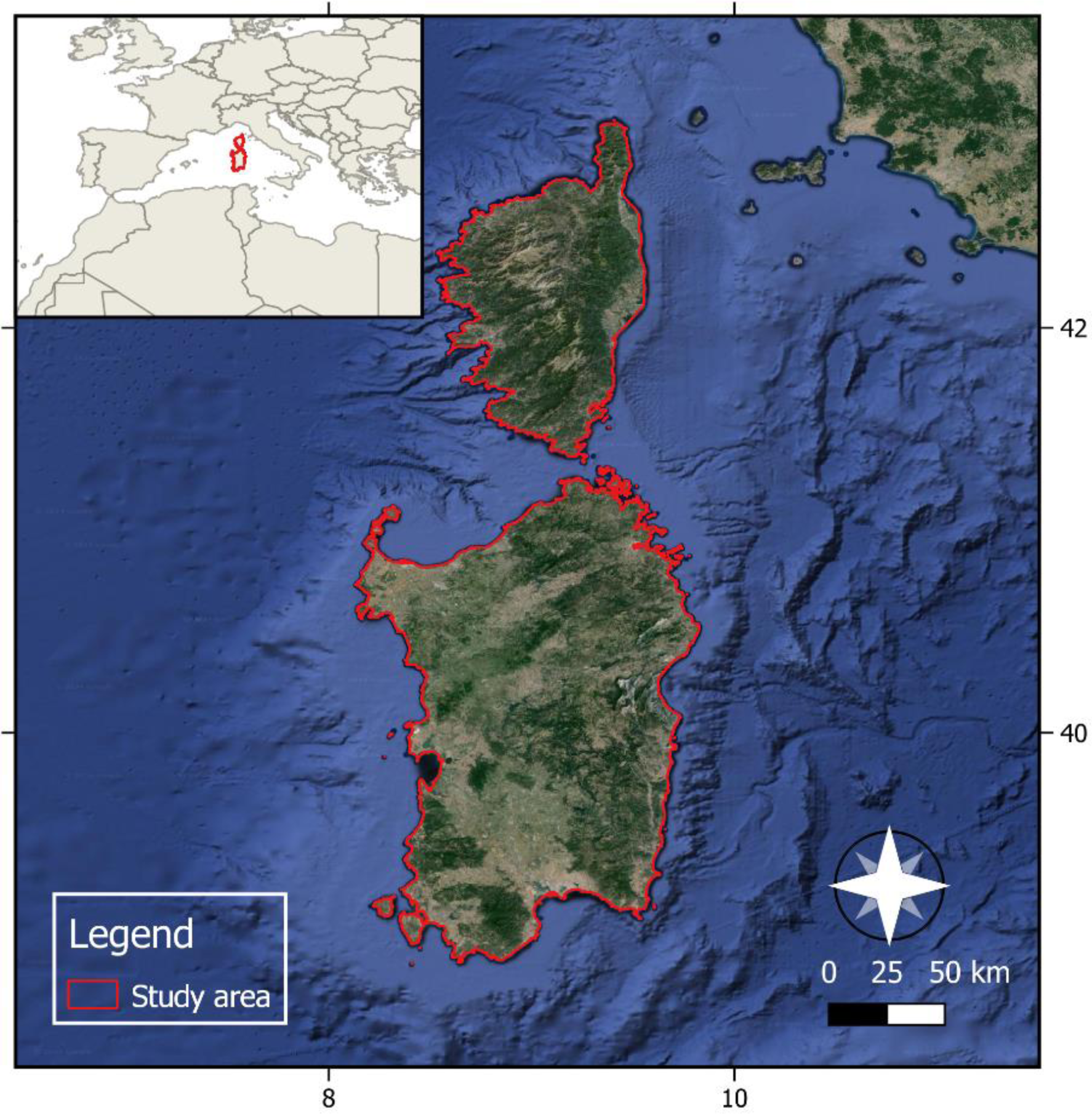
Study area. Satellite view of the Islands of Corsica and Sardinia, in the Northwestern Mediterranean Sea.

## 2. MATERIALS AND METHODS

### 2.1 Study area and taxa

Our study area is the Corsica-Sardinia microplate in the North-western Mediterranean (Fig: 1). Sardinia (24100 km^2^) is mainly mountainous (max altitude 1835 m) and hilly with the two main plains located on the western portion of the island. The climate is Mediterranean characterized by dry summers, mild winters and warm summers on the coasts and by cool summers on the mountains. Corsica (8722 km^2^) is characterized by a mountainous inland (max altitude 2700 m) and coastal plains. The climate in coastal regions is hot-summer Mediterranean and Warm-summer Mediterranean. At the highest elevation, there are small areas with an alpine climate. We focused our analyses on herpetofauna endemic (present only in one or both islands) and sub-endemic (being also present in few other localities) to the Corsica-Sardinia microplate (Table 1).

**Table 1.** Amphibians and reptiles considered in the study to investigate the effect of considering land use in fine scale SDMs future projections. The endemism column is divided as follows: E = endemic to Sardinia-Corsica microplate, S = Sub-endemic to Sardinia-Corsica microplate (i.e., being also present in few other localities). The “Occurrences” column refers to the number of occurrences used to train the model at the three selected resolutions.

| Species | Class | Endemism | Occurrences |  |  |
| --- | --- | --- | --- | --- | --- |
|  |  |  | 1 km<sup>2</sup> | 5 km<sup>2</sup> | 10 km<sup>2</sup> |
| *Algyroides fitzingeri* | Reptilia | S | 55 | 50 | 44 |
| *Archaeolacerta bedriagae* | Reptilia | S | 324 | 168 | 85 |
| *Chalcides ocellatus tiligugu* | Reptilia | S | 71 | 63 | 57 |
| *Discoglossus sardus* | Amphibia | S | 168 | 125 | 102 |
| *Euleptes europaea* | Reptilia | S | 44 | 37 | 30 |
| *Euproctus montanus* | Amphibia | E | 228 | 112 | 64 |
| *Euproctus platycephalus* | Amphibia | E | 27 | 25 | 22 |
| *Hyla sarda* | Amphibia | S | 276 | 210 | 157 |
| *Podarcis tiliguerta* | Reptilia | S | 1250 | 595 | 311 |
| *Salamandra corsica* | Amphibia | E | 254 | 117 | 54 |

### 2.2 Occurrences and background data

We obtained the occurrence data from two main sources: GBIF (Global Biodiversity Information Facility, https://www.gbif.org/) and iNaturalist (https://www.inaturalist.org/). We considered only occurrences with positional accuracy ≤ 1 km, and s not flagged as uncertain (Zizka et al., 2019). Moreover, to have a better correspondence with the land use and climate data, we retained in the analysis only the occurrences sampled after the year 2000. To reduce the effects of spatial autocorrelation, which can affect model performance (Dormann et al., 2007), we spatially filtered the observations, maintaining just one occurrence per cell and obtaining three different sets of filtered points per species, one for each resolution (1, 5, and 10 km^2^). The number of occurrence points used for model calibration ranged from 22 to 1250 (Table 1).

To minimize sampling bias in the occurrence data we used a target group approach for background points (Hortal et al., 2015; Garcia-Rosello et al., 2023), by using all occurrences of amphibians and reptiles collected in the study area. The assumption is that both species occurrences and background points should share approximately the same sampling bias (Ranc et al., 2017; Barber et al., 2022. We cleaned and spatially filtered background points following the same procedure described for the occurrence data. The number of background points used for model calibration was 3194 at 1 km^2^ resolution, 1140 at 5 km^2^ resolution, and 484 at 10 km^2^ resolution.

### 2.3 Bioclimatic variables and land use/ land cover variables

To calibrate the SDMs, we considered bioclimatic variables and land use/ land cover variables (hereafter “LULC” variables) (Table 2) at three different resolutions (1 km^2^, 5 km^2^, 10 km^2^). We obtained Land use variables from Copernicus (Harper et al., 2022) for 2020 and land use/land cover projections for 2050 from ESRI (https://livingatlas.arcgis.com/landcover-2050/). These projections were developed to be aligned with the (SSP 2-4.5) scenario. We then reclassified the twenty-three LULC classes for the year 2020 and the nine classes for the 2050 projection present in the study area into the eight LULC classes indicated in Table 2. We resampled both the current and the future land cover projections from their original resolution (300 m) to the three selected resolutions (1 km^2^, 5 km^2^, 10 km^2^). Land Cover was thus expressed as a percentage of surface on a 1 km^2^, 5 km^2^, or 10 km^2^ pixel. Moreover, we considered a water recurrence variable (Pekel et al., 2016), which provides information concerning the inter-annual behavior of the water surface and captures the frequency with which water returned from year to year in the 1984-2021 time frame. We resampled it from its original resolution (30 m^2^) to the three selected resolutions (1 km^2^, 5 km^2^, 10 km^2^). We assumed that water recurrence remains unchanged over time. To limit multicollinearity, we performed a Variance Inflation Factor (VIF) analysis with a threshold of VIF = 2 at the three different resolutions, obtaining a final set of ten predictors (Table 2).

**Table 2.** Bioclimatic and land use /land cover (LULC) variables considered for modelling the species distribution of 5 amphibian and 5 reptile species in Corsica and Sardegna. The column “Retained” indicates whether the variable was used or excluded because of multicollinearity.

| Predictor name | Description | Class | Retained |
| --- | --- | --- | --- |
| **Bio1** | Annual mean temperature | bioclimatic | NO |
| **Bio2** | Mean diurnal temperature range | bioclimatic | YES |
| **Bio4** | Temperature seasonality | bioclimatic | YES |
| **Bio5** | Maximum temperature of warmest month | bioclimatic | NO |
| **Bio6** | Minimum temperature of coldest month | bioclimatic | NO |
| **Bio8** | Mean temperature of wettest quarter | bioclimatic | NO |
| **Bio9** | Mean temperature of driest quarter | bioclimatic | NO |
| **Bio10** | Mean temperature of warmest quarter | bioclimatic | NO |
| **Bio11** | Mean temperature of coldest quarter | bioclimatic | NO |
| **Bio12** | Annual precipitation amount | bioclimatic | NO |
| **Bio13** | Precipitation of wettest month | bioclimatic | YES |
| **Bio14** | Precipitation of direst month | bioclimatic | NO |
| **Bio15** | Precipitation seasonality | bioclimatic | YES |
| **Bio16** | Precipitation of wettest quarter | bioclimatic | NO |
| **Bio17** | Precipitation of driest quarter | bioclimatic | NO |
| **Bio18** | Precipitation of warmest quarter | bioclimatic | NO |
| **Bio19** | Precipitation of coldest quarter | bioclimatic | NO |
| **Agriculture** | Percentage of agricultural area | LULC | NO |
| **Deciduous forest** | Percentage of deciduous forest | LULC | YES |
| **Evergreen forest** | Percentage of evergreen forest | LULC | YES |
| **Grassland and Shrubland** | Percentage of grassland and/or shrubland | LULC | YES |
| **Wetland** | Percentage of wetland | LULC | YES |
| **Settlement** | Percentage of settlement | LULC | YES |
| **Sparse vegetation** | Percentage of sparse vegetation | LULC | YES |
| **Bare area** | Percentage of bare area | LULC | YES |
| **Water recurrence** | Percentage of water recurrence | LULC | YES |

We obtained climate for the current time frame (averaged from 1970 to 2000) and for 2050 (averaged from 2041 to 2060) from Worldclim version 2.1 (Fick & Hijmans 2017) at the three selected resolutions. We excluded from our selection two of the 19 available bioclimatic variables (isothermality and temperature annual range) since they are a combination of other bioclimatic variables. For the 2050 projection, we selected five different General Circulation Models (GCMs: IPSL-CM6A-LR, MIROC6, MPI-ESM1-2-HR, MRI-ESM2-0, UKESM1-0-LL) and one shared socio-economic pathway (SSP 2-4.5) of the CMIP6 scenarios.

### 2.4 Species Distribution Models

We modelled the distribution of the species with MaxEnt, a machine learning approach that can estimate the most uniform distribution (maximum entropy) of presence points compared to the background locations given the constraints derived from the environmental data (Phillips et al., 2006). MaxEnt has been shown to outperform other SDMs’ algorithms also in cases of few occurrence data (Hernandez et al., 2006; Wisz et al., 2008; Valavi et al., 2021). Since model performance is sensitive to model specification (e.g., Anderson et al., 2011; Elith et al., 2010; Warren et al., 2014), for each taxon we compared thirty different combinations of feature classes and regularization multipliers and selected the best combination using the sample size corrected Akaike Information Criteria (AICc, Burnham and Anderson 1992). All the analyses were performed in R (R core Team 2022) with the “ENMeval” package (Muscarella et al., 2014; Kass et al., 2021). We calibrated all the models at three different resolutions (1 km^2^, 5 km^2^, and 10 km^2^) and with two different sets of explanatory variables: only bioclimatic variables (hereafter “C” models) vs bioclimatic and LULC variables (hereafter “CL” models) totaling 6 models for each species (tot=60 models).

We used a 10-fold cross-validation to separate training and testing data, and we evaluated model discrimination capacity by calculating the area under the receiver operating characteristic curve (AUC) (Jiménez-Valverde, 2012). Moreover, we tested whether the models selected differed significantly from what would be expected by chance by comparing their AUC values against a null distribution of 100 expected AUC values based on random data (Raes & ter Steege 2007; Bohl et al., 2019). For each model we assessed the variable importance as percent contribution (Phillips 2006). Lastly, we obtained current and future continuous suitability maps by projecting model results under current and future conditions. We binarized all models using the 10th percentile probability value measured with training occurrences, allowing for a 10% of omission errors (Brito et al., 2008). We evaluated the differences between the two modelling approaches (climate only vs climate and land use) at the three selected resolutions by comparing 1) models’ predictive performance in terms of AUC values, 2) changes in variable importance, 3) changes in the predicted suitability (in terms of km^2^ of suitable range predicted) at different time frames, 4) disagreement in spatial prediction (i.e., km^2^ for which the two different modelling approaches predicted a different outcome).

## 3. RESULTS

### 3.1 Evaluation of predictive performance

All SDMs gave AUC values significantly different than random except for the 10 km^2^ CL models of *Euproctus platycephalus* and *Podarcis tiliguerta.* In these two cases, the AUC score of 0.60 was not significantly higher than random and thus the two models were not further considered.

The average AUC values for C models were 0.73±0.1, 0.71±0.1, and 0.71±0.1 respectively for the 1 km^2^, 5 km^2^, and 10 km^2^ resolutions, while the average AUC values for CL models were 0.72±0.1 for all the three resolutions. At 1 km^2^ resolution, the inclusion of LULC variables improved predictive power for 10% (n=1) of the species (*Hyla sarda*), worsened it for 20% (n=2), and did not affect it for 70% (n=7) of the species. At 5 km^2^ resolution, the introduction of LULC variables improved predictive power for 20% (n=2) of the species, worsened it for 0% (n=0), and did not affect it for 80% (n=8) of the species. At 10 km^2^ resolution, the introduction of LULC variables improved predictive power for 20% (n=2) of the species, worsened it for 20% (n=2), and did not affect it for 60% (n=6) of the species.

### 3.2 Variable importance

In general, the addition of LULC variables affected the importance of bioclimatic variables in CL models in 3 non-exclusive ways, Table 3). Irrespective of the spatial resolution, LULC variables had higher and/or similar importance to bioclimatic variables in the model for ≥ 80% (n=8) of the species. For 20% to 30% of the species (depending on resolution) LULC variables changed the rank of the CLIM variables. For 10% to 25% of the species (depending on resolution LULC variables changed the relative importance, but not the ranking, of the bioclimatic variables.

**Table 3.**
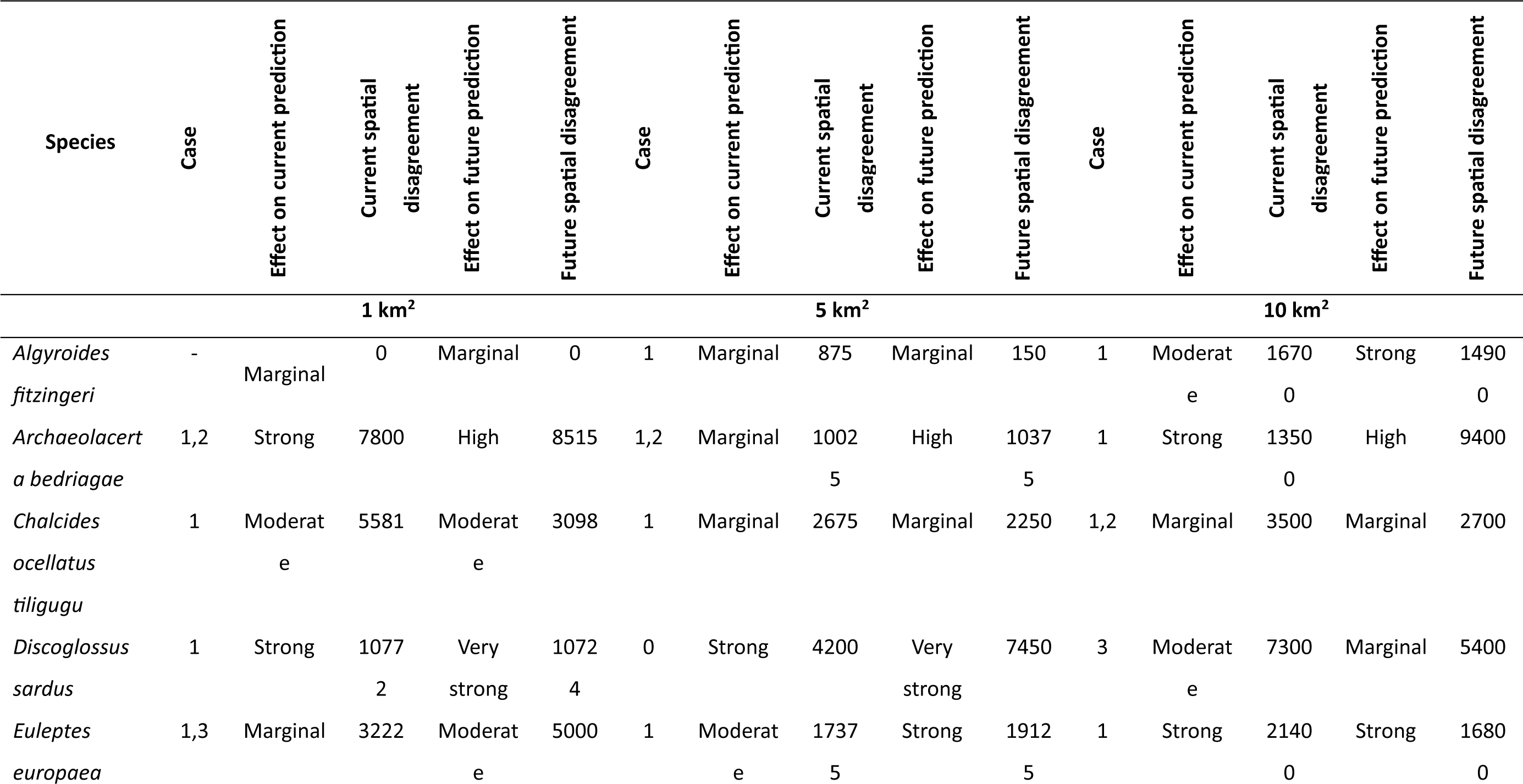
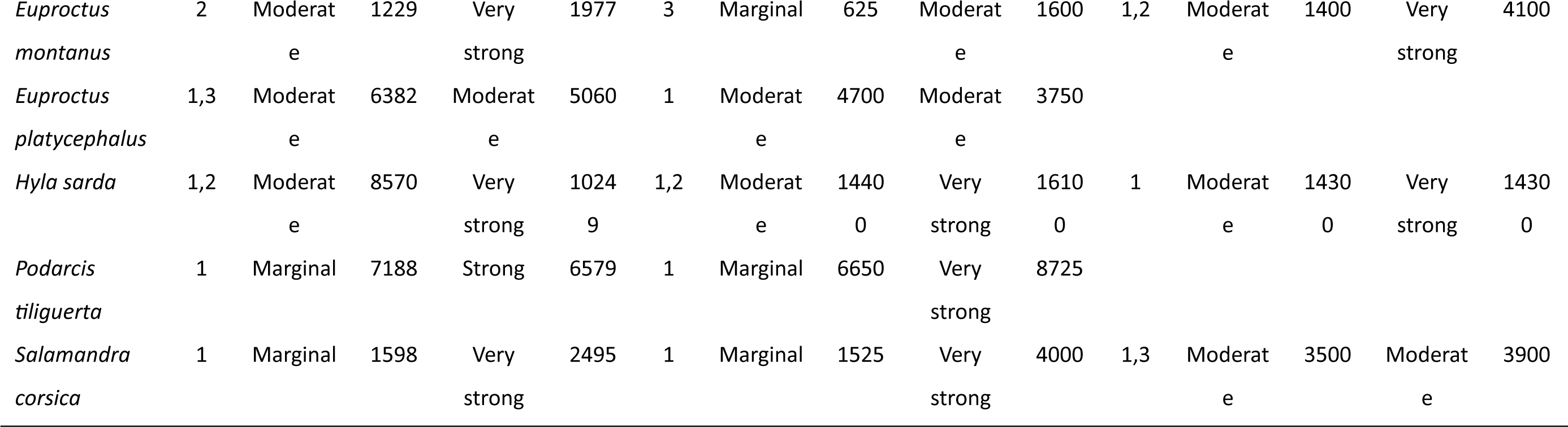
Effects of land use/land cover variables on variable importance in climate land use models (CL) compared to bioclimatic models. The Case column indicates the effect of the introduction of land use/ land cover variables (LULC). Case 1: LULC variables had higher/similar importance to bioclimatic variables in the model. Case 2: LULC variables changed the order of importance of the CLIM variables, Case 3: LULC variables changed the relative importance (but not the order) of the bioclimatic variables, the symbol – indicates that the introduction of LULC variables had no effects on variable importance. The other columns indicate the effect of LULC variables on current and future predictions of suitable area (km^2^): Marginal = difference in suitable area ≤ 5%, Moderate: difference in suitable area ≤ 20%, Strong: difference in suitable area ≤ 40%, Very strong: difference in suitable area > 40%.

### 3.3 Model predictions

#### 3.3.1 Current predictions

The effect of including LULC variables in the SDMs on the model output was heavily dependent on the resolution and on the species being considered. In general, the disagreement between C and CL increased with coarser resolution, while remaining somewhat consistent within the same species (Table3, Fig. 2). For models at 1km^2^ resolution the mean spatial disagreement was 5,234 km^2^, ranging from 0 km^2^ (*Algyroides fitzingeri*) to >10,000 km^2^ (*Discoglossus sardus*). At this resolution, 40% of the species (n=4) showed a ≤5% difference in the amount of predicted suitable area between C and CL models while 40% (n=4) showed a 6-20% difference and 20% (n=2) showed a 21-40% difference (Table 3, Fig. 2).

**Figure 2.**
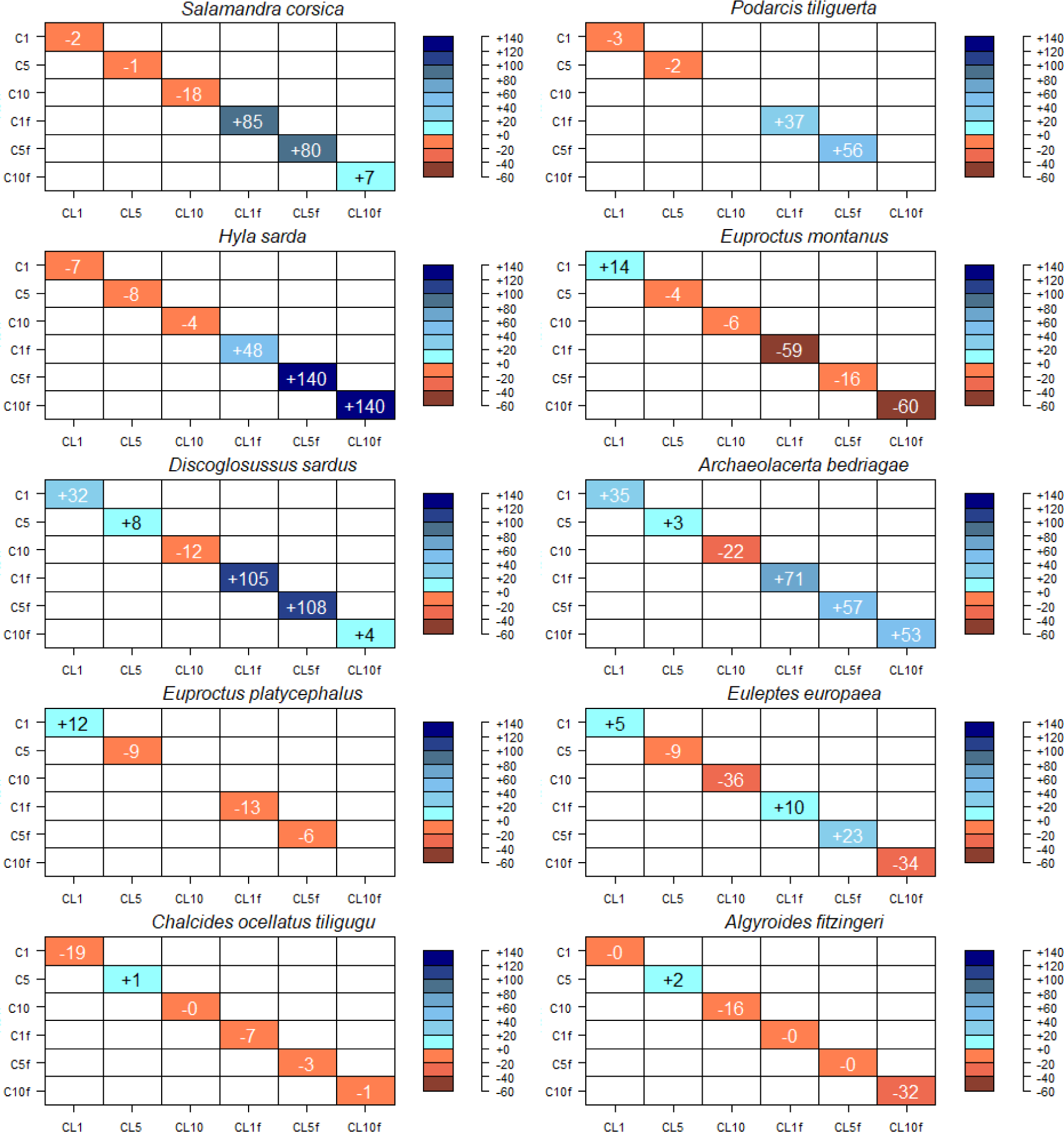
Differences in predicted suitable area (km^2^) at the current and future time frames between bioclimatic models (C for current and Cf for future) and models including bioclimatic and land use/land cover variables (CL for current and CLf for future) for the 10 species modelled and at three different resolutions (1, 5 and 10 km^2^). The result is expressed as a percent difference in the predicted suitable area (km^2^) between the models indicated by the columns of the matrix and those indicated by the row. The color scale indicates the sign of the difference (positive in blue and negative in red) and the magnitude (darkest colors indicated a larger difference).

For models at 5 km^2^ the mean spatial disagreement was 6,305 km^2^ ranging from 625 km^2^ (*Euproctus montanus*) to >17,000 km^2^(*Euleptes europaea*). At this resolution, 60% (n=6) of the species showed a ≤5% difference in the amount of predicted suitable area between C and CL models while 40% (n=4) showed a 6-20% difference (Table 3, Fig. 2).

For models at 10 km^2^ resolution the mean spatial disagreement was 10200 km^2^ ranging from 1,400 km^2^ (*Euproctus montanus*) to >21,000 km^2^ (*Euleptes europaea*). At this resolution, changes were more important, with only 25% (n=2) of the species showing a ≤5% difference in the amount of predicted suitable area between C and CL models, while 50% (n=4) of the species showed 6-20% change and 25% (n=2) showed a 21-40% difference (Table 3, Fig. 2).

#### 3.3.2 Future predictions

Considering future projections, all results followed the same patterns but with stronger differences. For models at 1km^2^ resolution the mean spatial disagreement was 5379 km^2^ ranging from 0 (*Algyroides fitzingeri*) to 10724 km^2^ (*Discoglossus sardus*). At this resolution 10% (n=1) of the species showed ≤5% difference in the amount of predicted suitable area between C and CL models, 30% (n=3) showed 6-20% differences, 10% (n=1) showed 21-40% differences and 50% (n=5) showed >40% differences (Table 3, Fig. 2).

For models at 5 km^2^ the mean spatial disagreement was 7352 km^2^ ranging from 150 (*Algyroides fitzingeri*) to 19125 km^2^ (*Euleptes europaea*). At this resolution 10% (n=1) of the species showed ≤5% difference in the amount of predicted suitable area between C and CL models, 30% (n=3) showed 6-20% differences, 10% (n=1) showed 21-40% differences and 50% (n=5) showed >40% differences (Table 3, Fig. 2).

For models at 10 km^2^ resolution the mean spatial disagreement was 8,938 km2, ranging from 2,700 (*Chalcides ocellatus tiligugu*) to 16,800 km^2^ (*Euleptes euroapea*). At this resolution 25% (n=2) of the species showed ≤5% difference in the amount of predicted suitable area between C and CL models, 10% (n=1) showed 6-20% differences, 25% (n=2) showed 21-40% differences and 40%(m=3) showed very strong differences (Table 3, Fig. 2).

#### 3.3.3 Predicted trends

Considering projected gain and losses in the distributions of all species, C models predicted a percent change in suitability ranging from -63% to +31% at 1km^2^ resolution, from -81% to +37% at 5 km^2^ resolution, and from -85% to +17% at 10 km^2^ resolution (Figure 3). CL models predicted trends ranging from -85% to +43% at 1 km^2^ resolution, from -59 to +41% at 5km^2^ resolution, and from -84% to -7% at 10 km^2^ resolution (Figure 4).

**Figure 3.**
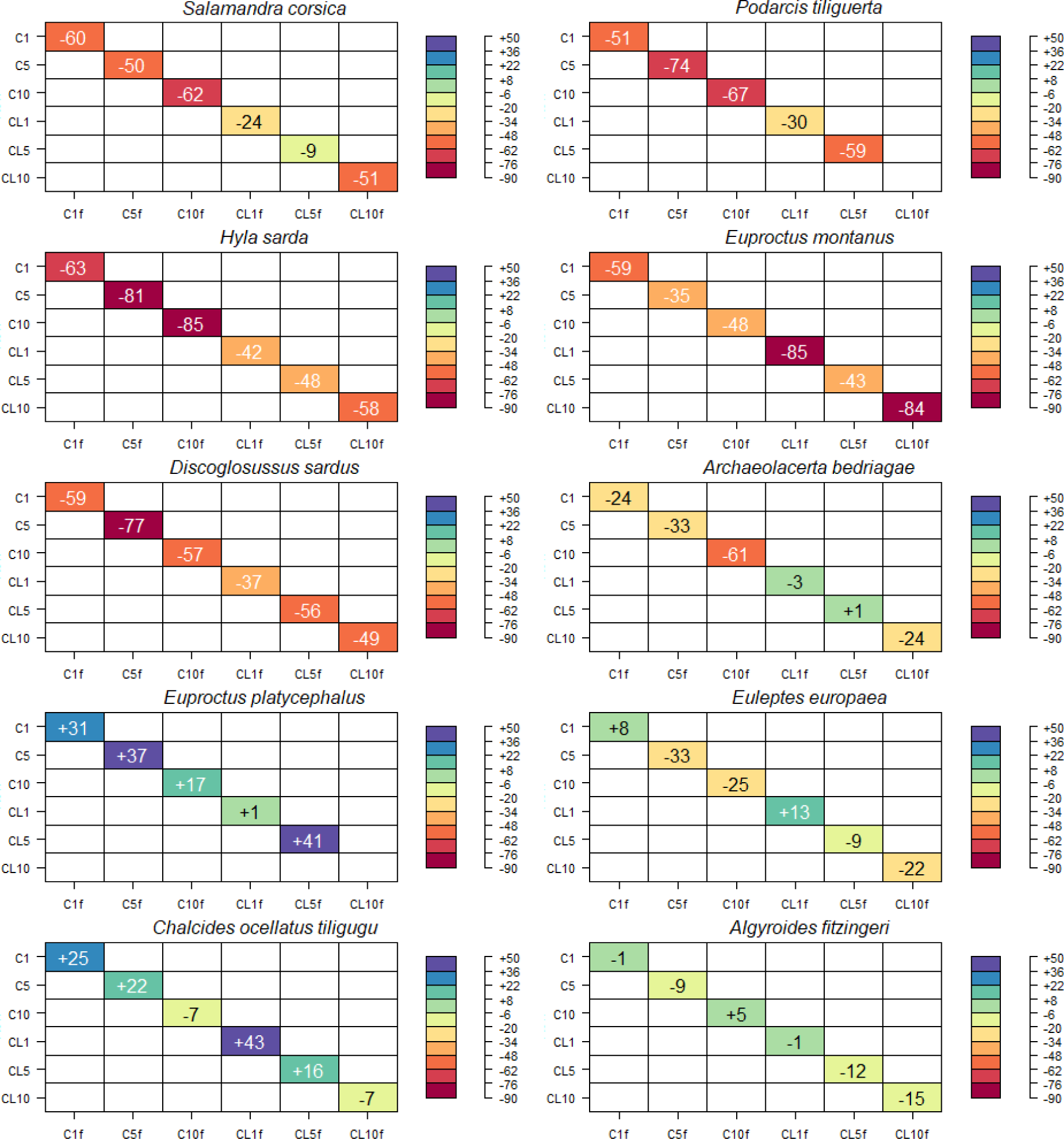
Trends in the suitable area (km^2^) at different resolutions (1, 5 and 10 km^2^) predicted by bioclimatic models (C *for current and Cf for future*) and models including bioclimatic and land use/land cover variables (CL *for current and CLf for future*) for the ten species modelled. The result is expressed as a percent difference between future and current predictions of suitable area. Warm colors indicate a predicted future loss in a suitable area and the intensity of the color increases with the magnitude of the loss. Cold colors indicate a predicted future gain in suitable area and the intensity of the color increases with the magnitude of the gain.

**Figure 4.**
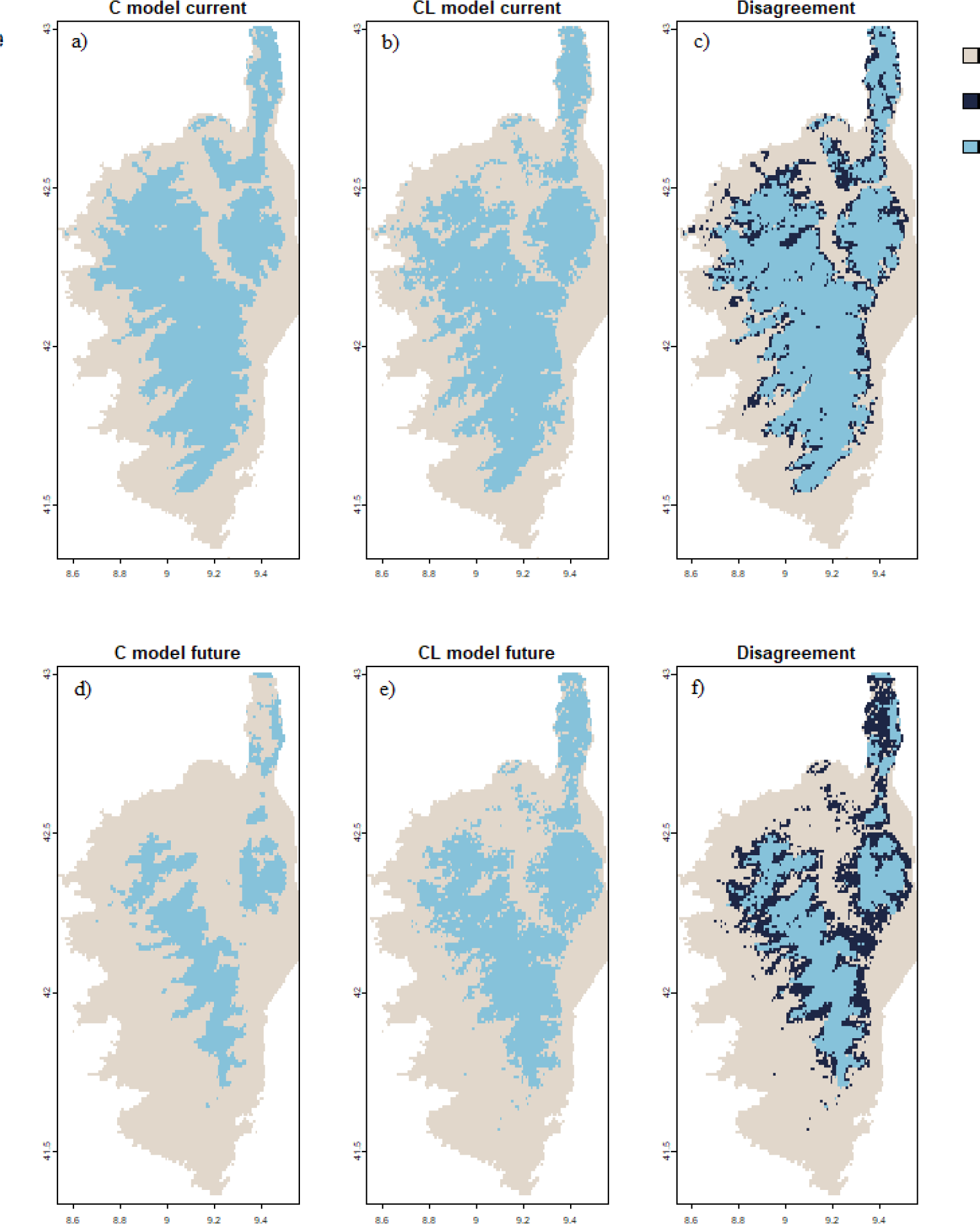
Suitability maps at 1km^2^ resolution for *Salamandra Corsica*. First row: maps of current suitability obtained with C models (a) and CL models (b) and disagreement between the two (c). Second row: maps of future suitability obtained with C models (d) and CL models (e) and disagreement between the two (f).

Overall, for 5 species (*Salamandra corsica*, *Hyla sarda*, *Discoglossus sardus*, *Podarcis tiliguerta*, *Archaeolacerta bedriagae*) we measured a consistent change in the trends when LULC variables are considered in the SDMs. In particular the CL models, when compared to the C models, always predicted a smaller decrease (mean across species =-30%, min=-18% for *Podarcis tiliguerta*, max=-67% for *Archeolacerta bedriagae*) in suitability, irrespective of the spatial resolution considered. At the opposite, for the *Euproctus montanus* the inclusion of LULC variables in the model produced a greater decrease in suitability (min=+8 at 5 km^2^ resolution, max=+36% at 10 km^2^ resolution) for all spatial resolutions.

For the remnant 4 species (*Chalcides ocellatus tiligugu*, *Euproctus platycephalus*, *Euleptes europaea*, *Algyrodes fitzingeri*) the effect of including LULC variables on the predicted trends in suitability did not show a clear pattern, always depending on the resolution considered.

## DISCUSSION

Bioclimatic species distribution models are possibly the most widely used tool in global change biology, biogeography, and conservation to project the response of species to global changes. When considering regional to continental spatial scales, most of the models are calibrated considering only bioclimatic variables (e.g., Maiorano et al. 2013) under the assumption that other factors influencing species distribution have only a marginal importance. Land use variables are a clear example: often used for smaller areas in higher resolution studies as proxies for habitats, these variables are often neglected at larger spatial scales assuming that they are not important in shaping species distribution. Thuiller et al. (2004) clearly demonstrated that at the scale of Europe and with a resolution of 50 km^2^ the addition of land use variables to model the potential distribution of herpetofauna has no influence on the predictive power of SDMs.

Here we explored the same issue but considering a study area that has a clear biogeographical delineation, sharp geographical boundaries, and a fauna rich in endemic species and subspecies in both reptiles and amphibians. We focused our research questions not only on the predictive power of the models (i.e., their AUC) but also on the geographical output (the maps of potential distribution), and on the explanatory component (changes in variable importance). Considering a set of 10 species, we calibrated two SDMs for each species, one including climate only variables, and the other including the same set of climate variables plus land use.

We found that the inclusion of land cover can profoundly change the models’ output. Consistently with previous studies (Thuiller et al., 2004), the inclusion of LULC variables did not change the models’ AUC for 80% of the species at all resolutions. Although all considered models achieved an AUC that was better than that of a null model (except for the models for *Podarcis Tiliguerta* and for *Euproctus plathycephalus* at 10 km^2^ resolution), for some of the considered species (*Discoglossus sardus, Hyla sarda, Algyroides fitzingeri, Euleptes europaea, Podarcis tiliguerta*) prediction success was poor (0.6 < AUC ≤ 0.7; Araujo et al., 2005) at all resolutions. This was likely to be a result of the ubiquity of many of the species together with the restricted calibration area, that caused the occurrence points to capture much of the environmental variation used for model calibration (Tingley et al., 2009; Luoto et al., 2005).

Considering the explanatory power of the SDMs, land use variables constantly changed the rankings or the values of variables importance. Only one taxon, the Fitzinger’s algyroides (*Algyroides fitzingeri*) at the resolution of 1 km^2^ retained the same variable importance for both SDMs. For all other taxa the introduction of LULC variables influenced variable importance at all resolutions. In most of the cases (≥80%), when LULC variables were included, they represented the three most important variables, and for 30% of the species LULC variables affected also the rank of importance of the climatic variables and/or changed their relative importance. Contrary to what expected with previous studies (Thuiller et al., 2004; Luoto et al., 2007), the rank of importance of LULC variables did not increase with higher spatial resolutions.

Considering the projections of the models over the study area under current environment, the difference between climate only SDMs and climate plus land use SDMs was overall limited at all spatial resolutions (mean=10%, min=0%, max=36%). With few exceptions, CL models tended to predict the same or slightly less suitable surface than C models (Fig. 2). This result agrees with previous studies and can be explained by the fact that climate generally varies on a larger scale than land cover and as a result C models often produced smoother, more continuous predictions than CL models (Tingley e al., 2009). This result also supports the notion that land cover may be able to refine the predictions of C models by identifying areas that are climatically suitable, but that are inhospitable owing to the effects of habitat loss and degradation (Pearson et al., 2004).

When considering projections to future environment, the difference in amount of predicted suitable area between the two approaches is much greater and a pattern emerges. For the species that are predicted to face extensive range contraction (> 50%) according to C models (4 amphibians, 2 reptiles), the difference is large (mean= 69%, min=4%, max=167%) and this result is consistent for all resolutions. For species that are predicted not to face extensive range contraction by C models (i.e., thee species, 3 reptiles and 1 amphibian that are predicted either to lose a moderate portion of their range, or maintain it or expand it) the difference between the two approaches is smaller (mean= 11%, min=0%, max=34%) and varies with resolution, without a clear pattern.

For the first group of species (*Salamandra corsica, Hyla sarda, Euproctus montanus, Discoglossus sardus, Podarcis tiliguerta, Archaeolacerta bedriagae*) C and CL models project changes in suitability of different entity. For all these species except one (*Euproctus montanus*), in the future scenario, C models tend to predict considerably less (mean = -72%, min= -4%, max=-167%) suitable surface compared to CL models. When looking at variable importance the difference is attributable to the fact that when introducing LULC variables they mainly are among the 3 most important explanatory variables and/or change the relative importance of bioclimatic variables. Ecologically, the use of LULC variables seem to identify areas that would be unsuitable based on the climate only but qualify as suitable when also considering the land cover, likely identifying important local resource such as cover (Seoane et al., 2004). One example of that is represented by *Salamandra corsica*, which is known to depend on densely vegetated and topographically complex habitats, which reduce temperature fluctuations (Escoriza & Hernandez 2021). For this species C models predicted a range loss of -60%, -50% and -62% respectively at 1, 5 and 10 km^2^ resolution due mainly to its negative response to a change in precipitation seasonality (bio15) and mean diurnal temperature range (bio2) values. In CL models LULC variables (i.e., evergreen forest and sparse vegetation) were between the first 3 variables and the predictions identified several additional suitable areas in future projections (Fig. 4). There is one species in this group for which the pattern is opposite: *Euproctus montanus*. For this species CL models predict a more dramatic decrease in suitability (-85%, -43%, -84% respectively at 1, 5 and 10 km^2^ resolution) than C models (-59%, -35%, 48% respectively at 1, 5 and 10 km^2^ resolution, Fig. 5). For this species, LULC variables were not among the 3 most important, but they affected the order of importance of the bioclimatic variables. In this case precipitation seasonality (bio15) was the most important variable in the C models, while temperature seasonality (bio4) was the most important variable in the CL model. These results show that for these species considering or not LULC variables produces different conclusions in term of future species conservation status. Since validating future model projections is not possible (Elith et al., 2010), our results outline the importance of closely considering the ecology of each species avoiding, as much as possible, approaches that treat all species at the same way.

**Figure 5.**
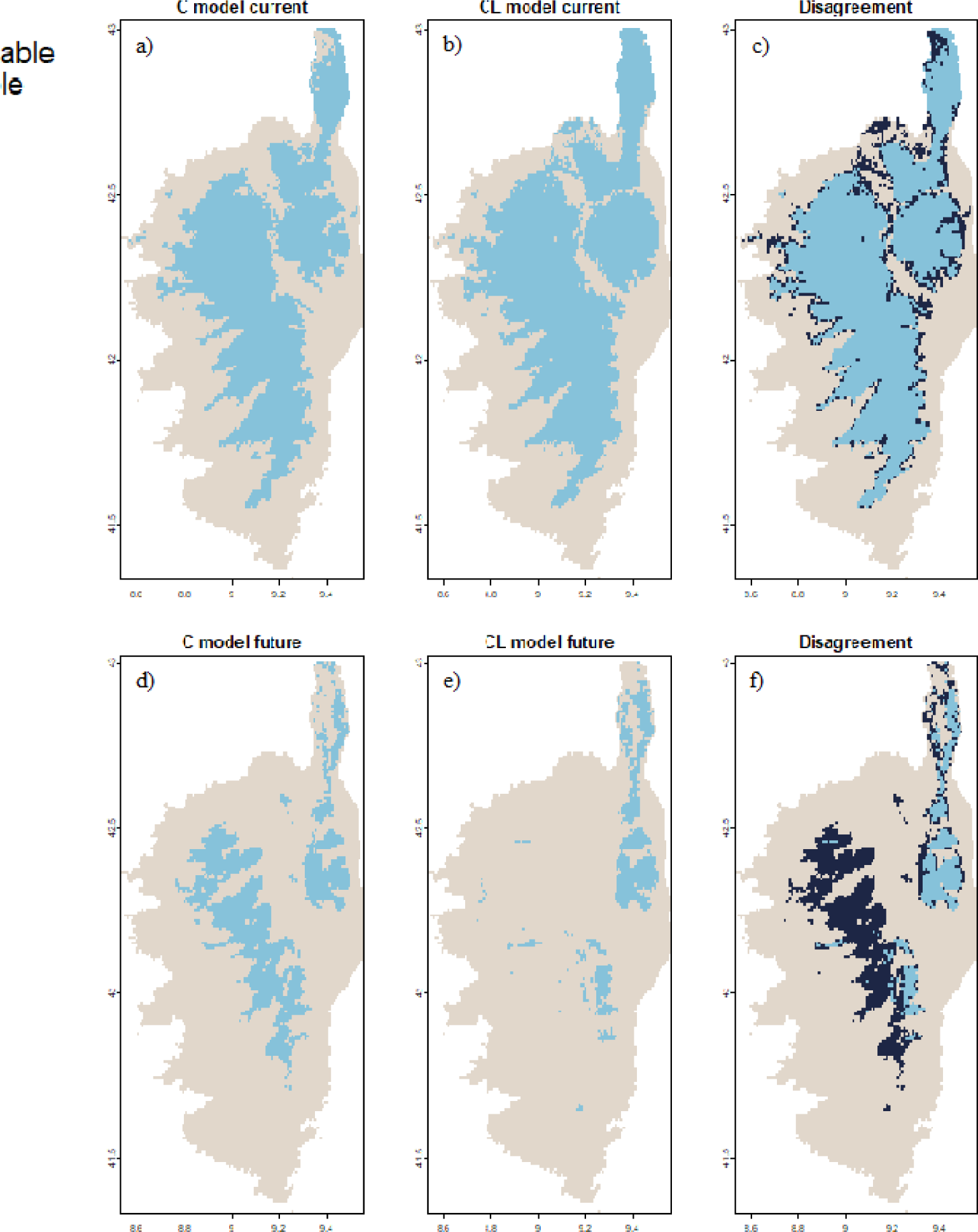
Suitability maps for *Euproctus montanus*. First row: maps of current suitability obtained with C models (a) and CL models (b) and disagreement between the two (c). Second row: maps of future suitability obtained with C models (d) and CL models (e) and disagreement between the two (f).

For the second group of species (*Euproctus plahycephalus, Chalcides ocellatus tiligugu, Euleptes europaea* and *Algyroides fitzingeri*) C models predicted mainly a moderate range change (mean=5.9%, min=-6%, max=32%). For some of these species the introduction of variables only moderately affected the amount of future suitability predicted (e.g., *Euproctus platycephalus* and *Chalcides ocellatus tiligugu*), for others C models predicted the same, slightly less or slightly more suitable surface depending on the considered resolutions (e.g., *Euleptes europaea* and *Algyroides fitzingeri*). In terms of predicted range change, the difference between the two approaches did not show any consistent trend.

The two approaches did not differ only concerning the predicted suitable surface but also in terms of the location of suitable areas. The disagreement varied across species, but it was generally high for both the current and future scenarios and increased with decreasing resolutions (Table 3). Interestingly we observed high disagreement also for species for which the amount of suitable area predicted was similar (e.g, *Podarcis tiliguerta*, Fig. 6).

**Figure 6.**
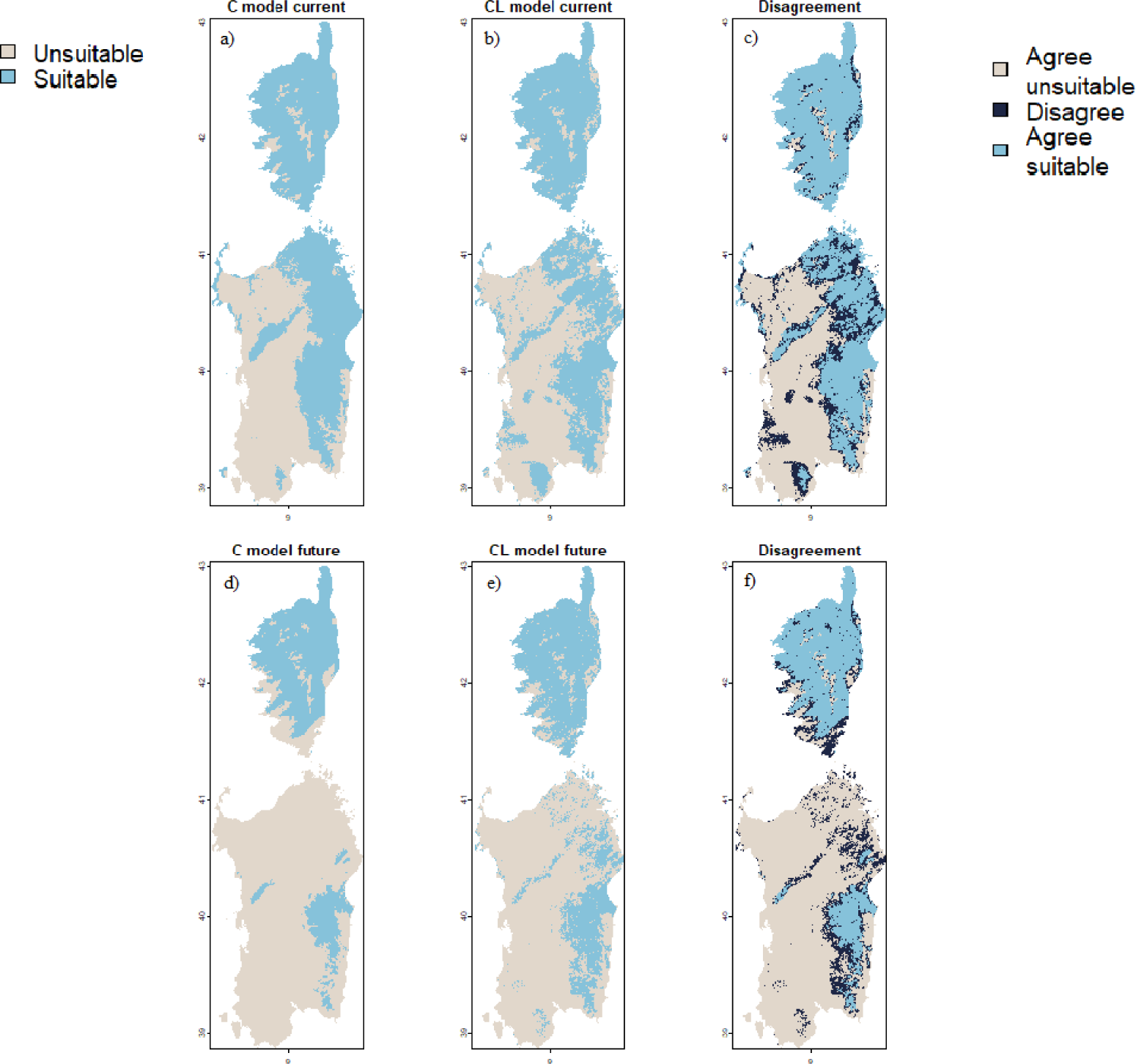
Suitability maps for *Podarcis tiliguerta*. First row: maps of current suitability obtained with C models (a) and CL models (b) and disagreement between the two (c). Second row: maps of future suitability obtained with C models (d) and CL models (e) and disagreement between the two (f).

Predicting potential future range shift in biodiversity is fundamental to identify priority areas for conservation (e.g., climatic refugia, Cavalcante et al., 2024), but it requires to include the main shift drivers in the modelling exercise. Our results point out that the implications of non-including land use change variables are non-negligible and should be considered on a taxon-specific basis. From our results we could not derive clear indications regarding if land use variables should be included or not. For several species considering climate change only could be more conservative (i.e. representing the worst-case scenario of range contraction) but for other species the opposite is true. Moreover, since the two approaches have considerable predictive disagreement, a cautious approach would compare both modelling approaches and focus on the agreement areas.

